# Extraction of Protein Dynamics Information Hidden in Cryogenic Electron Microscopy Maps Using Deep Learning

**DOI:** 10.1101/2020.02.17.951863

**Authors:** Shigeyuki Matsumoto, Shoichi Ishida, Mitsugu Araki, Takayuki Kato, Kei Terayama, Yasushi Okuno

## Abstract

Technical breakthroughs in cryogenic electron microscopy (cryo-EM)-based single-particle analysis have enabled the structures of numerous proteins to be solved at atomic or near-atomic resolutions, including extremely large macromolecules whose structures could not be solved by conventional techniques. Determining the dynamics properties of these macromolecules, based on their solved structures, can further improve our understanding of their functional mechanisms. However, such analysis is often hampered by the large molecular size and complex structural assembly, making both experimental and computational approaches to determine dynamics properties challenging. Here, we report a deep learning-based approach, DEFMap, to extract the dynamics information “hidden” in cryo-EM density maps. By relying only on cryo-EM maps, DEFMap successfully provided dynamics information equivalent to that determined from molecular dynamics (MD) simulations and experimental approaches at the atomic and residue levels. Additionally, DEFMap could detect dynamics changes associated with molecular recognition and the accompanying allosteric conformational stabilizations, which trigger various biological events such as signal transduction and enzyme catalysis. This approach will provide new insights into the functional mechanisms of biological molecules, accelerating modern molecular biology researches. Furthermore, this advanced strategy combining experimental data, deep learning approaches, and MD simulations would open a new multidisciplinary science area

## Main

The functions of a protein under physiological conditions are associated with its three-dimensional (3D) structure and dynamic behavior in solution. High-resolution investigations of the structural and dynamics properties of proteins are, therefore, essential for elucidating the molecular mechanisms underlying their biological functions, such as the regulation of cellular signaling mediated by protein–protein interactions and metabolic reactions catalyzed by enzymes^1,2^. Various techniques have been developed to determine protein structures, such as X-ray crystallography, nuclear magnetic resonance (NMR), and cryogenic electron microscopy (cryo-EM) single-particle analysis (SPA)^3–6^. Dynamics information has also been quantitatively obtained through several experimental and computational approaches, such as NMR, hydrogen-deuterium exchange (HDX) mass spectrometry (MS)^7^, and molecular dynamics (MD) simulations^8^.

Recent breakthroughs in cryo-EM SPA^3,9^ have enabled the determination of the structures of numerous biological molecules at atomic or near-atomic resolution^10,11^, including extremely large and complex macromolecules^12–15^ that cannot be solved using conventional techniques. However, investigating the dynamics of such molecules is technically challenging due to the large molecular sizes and complex structural assemblies involved.

To address these challenges, we developed a deep learning-based approach, named as “dynamics extraction from cryo-EM map” (DEFMap), to obtain dynamics information of proteins using a cryo-EM map alone (Fig. 1a and Supplementary Figure 1). The 3D cryo-EM maps solved by SPA are reconstructed from a large number of two-dimensional (2D) images of molecular particles identified in a micrograph^3–5^. Specimens used for cryo-EM SPA are prepared by rapidly freezing a solution in which the proteins adopt variable conformations. Thus, it has been generally known that the local map intensities in reconstructed 3D cryo-EM maps tend to correlate with dynamics properties, i.e., weaker intensities correspond to more flexible regions of the macromolecule. Therefore, their dynamics properties could be “hidden” in the reconstructed cryo-EM maps. However, no strategies for quantitatively extracting the hidden dynamics information directly from 3D cryo-EM maps alone have been reported.

**Fig. 1:**
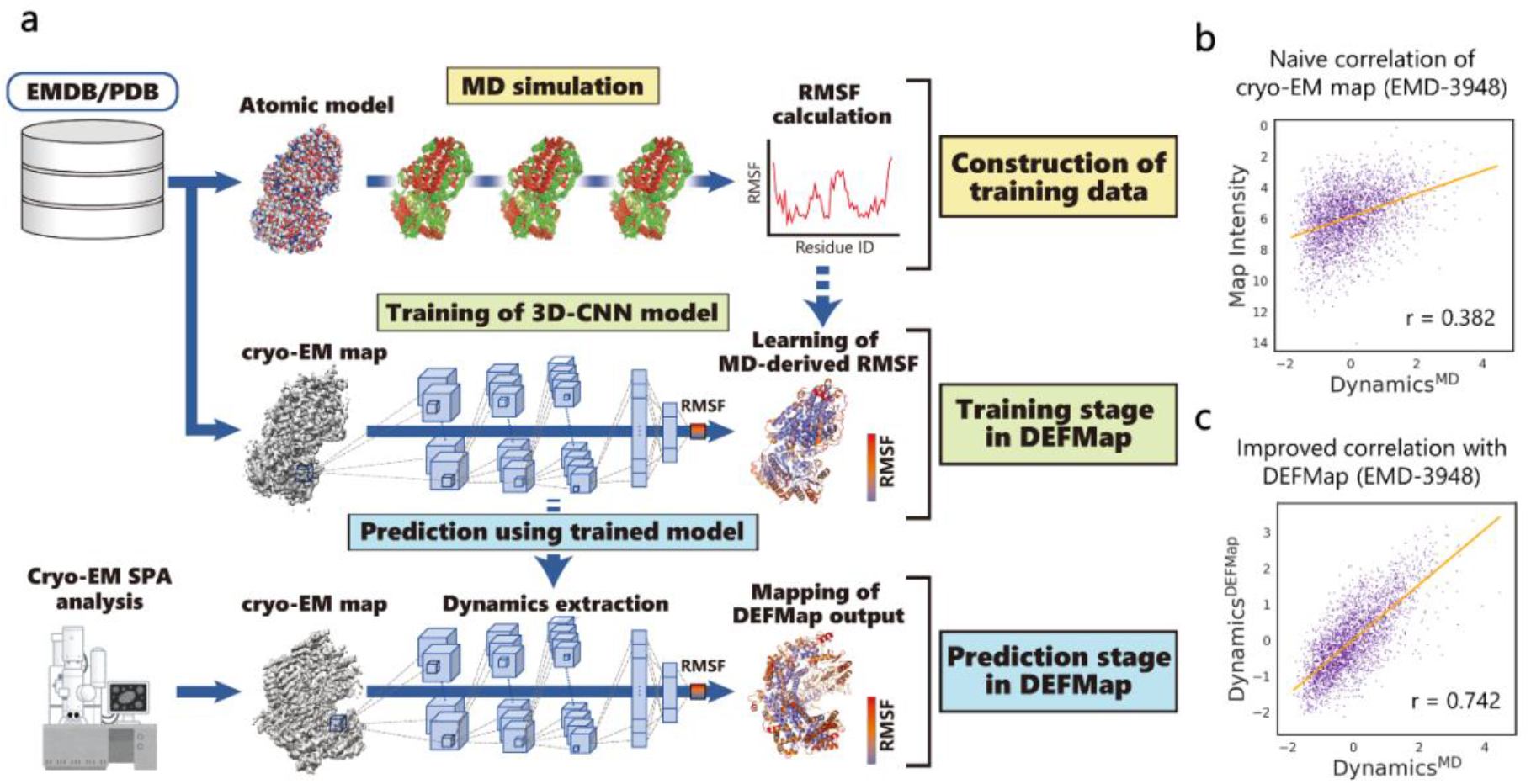
DEFMap-based extraction of dynamics features from cryo-EM maps. **a,** Whole workflow of DEFMap. Model training is carried out using EMDB/PDB-derived macromolecules which are conveniently handled in all-atom MD simulations. In the stage of training data-construction, the dynamics properties are derived from RMSF values (for heavy atoms) calculated from MD simulations. In the training stage, a 3D-CNN model in DEFMap (Supplementary Figure 1) learns the relationship between dynamics values and density data at the corresponding positions. In the prediction stage, for other cryo-EM maps not included in the training dataset, the trained model predicts dynamics values based on input density data. In this study, 25 macromolecules were used to validate and train the DEFMap model, and other 9 macromolecules were used for dynamics predictions using the trained model and further structural analyses. **b,** A correlation plot between the cryo-EM map intensities and Dynamicsmd for EMD-3948 (entry 1 in Supplementary Table 2). Dynamicsmd were calculated from RMSF values derived from MD simulations. **c,** A correlation plot between Dynamicsdefmap and Dynamicsmd for EMD-3984 (entry 1 in Supplementary Table 2). *r* denotes the correlation coefficient.

To quantitatively extract the hidden dynamics information at the atomic or residue level only from density data in the cryo-EM map, DEFMap is constructed by using a deep learning method and MD simulation (Fig. 1a). The deep learning method is designed to learn the relationship between local density data and dynamics information. Although the quantitative dynamics information is required to train the neural network model, it is not realistic to obtain enough dynamics information from existing experimental methods. To overcome this limitation, we performed MD simulations using cryo-EM map-derived atomic structures in the Protein Data Bank (PDB)^16^, and calculated the root-mean-square fluctuation (RMSF) representing atomic fluctuations as dynamics information (see the stage of training data-construction in Fig. 1a). MD simulations have been widely used to elucidate the detailed dynamic behavior of biological molecules^8,17,18^, as MD techniques and computing performance have advanced. Next, we trained the 3D-convolutional neural network (3D-CNN)^19–21^ model to learn the relationship between cryo-EM density data and MD-derived RMSF values, to capture 3D patterns of the cryo-EM density data reflecting protein dynamics (see the training stage in Fig. 1a). 3D-CNN has been widely used to detect or classify patterns in 3D objects in various fields^22–24^, and has exhibited remarkable performance in the processing of 3D cryo-EM maps^25–28^. Finally, using the trained 3D-CNN model, DEFMap can directly and quantitatively extract “hidden” dynamics information as RMSF values only from 3D cryo-EM density data of a new target protein (see the prediction stage in Fig. 1a). Note that at the prediction stage, DEFMap does not require any MD calculations. For visualization, DEFMap maps residue-specific RMSF values averaged over each residue after normalization (termed Dynamicsdefmap) onto the corresponding atomic models.

At first, to evaluate the performance of DEFMap, we collected 25 cryo-EM maps from the Electron Microscopy Data Bank (EMDB)^29^ (Supplementary Table 1) and used them to train the DEFMap 3D-CNN model (see Methods section for the details of the training dataset, MD simulation and training). Within the dataset, the map intensities tended to correlate, to some extent, with the residue-specific RMSF values obtained from MD (termed Dynamicsmd) as shown in Fig. 1b and Supplementary Table 2. However, this correlation was insufficient to directly estimate the dynamics properties from the cryo-EM maps, as the average correlation coefficient *r* was only 0.464 ± 0.164. Using the same dataset, we performed leave-one-out cross-validation^30^ to accurately evaluate the performance of DEFMap within the dataset. The results showed an improved correlation (*r* = 0.665 ± 0.124) between the Dynamicsdefmap and Dynamicsmd outputs (Fig. 1c and Supplementary Table 2). This finding indicates that DEFMap can efficiently extract dynamics features from 3D cryo-EM density data.

We subsequently trained the 3D-CNN model of DEFMap using the full training dataset (25 cryo-EM maps) and then tested DEFMap on other three additional cryo-EM maps (EMDB/PDB entry; EMD-4241/6FE8^31^, EMD-7113/6BLY^32^, and EMD-20308/6PCV^33^) to further evaluate its potential for dynamics analysis (Supplementary Table 3). They were selected to exhibit distinct structural aspects in terms of secondary structure contents (α-helix/β-strand/others, see legend of Figure 2a). Particularly, experimental dynamics of EMD-20308/6PCV has been reported and is compared with Dynamicsdefmap later. The performance of DEFMap in extracting dynamics features from cryo-EM maps is illustrated in Figure 2. The dynamics values calculated by the trained DEFMap model showed good correlation with those derived from MD simulations at both the atomic (*r* = 0.704, 0.726 and 0.673, respectively) and residue (*r* = 0.727, 0.748 and 0.711, respectively) levels (Supplementary Fig. 2). The results showed that DEFMap could extract accurate dynamics information from the cryo-EM map data alone. Moreover, the mappings of Dynamicsdefmap onto the corresponding atomic models demonstrated that DEFMap captured conformational aspects such as rigidity in the protein interior and flexibility of the solvent-exposed secondary structure elements, with accuracy similar to that of MD simulations (Fig. 2a–c, upper panels). The Dynamicsdefmap data agreed well with the Dynamicsmd data throughout the protein (Fig. 2a–c, bottom panels). However, in some regions, the DEFMap-based calculations failed to extract accurate dynamics properties. We hypothesize that a low local map resolution hinders accurate extraction of dynamics data. In fact, lower overall map resolutions (particularly those >8 Å) resulted in inferior performances (Fig. 2d and Supplementary Fig. 3). This tendency reflects the loss of detailed structural information in low-resolution maps. The apparent resolution dependence of the DEFMap model suggests that its performance will be enhanced alongside the continuous improvements in resolution associated with the development of advanced equipment, such as cold field emission guns^34,35^, for cryo-EM data acquisition.

**Fig. 2:**
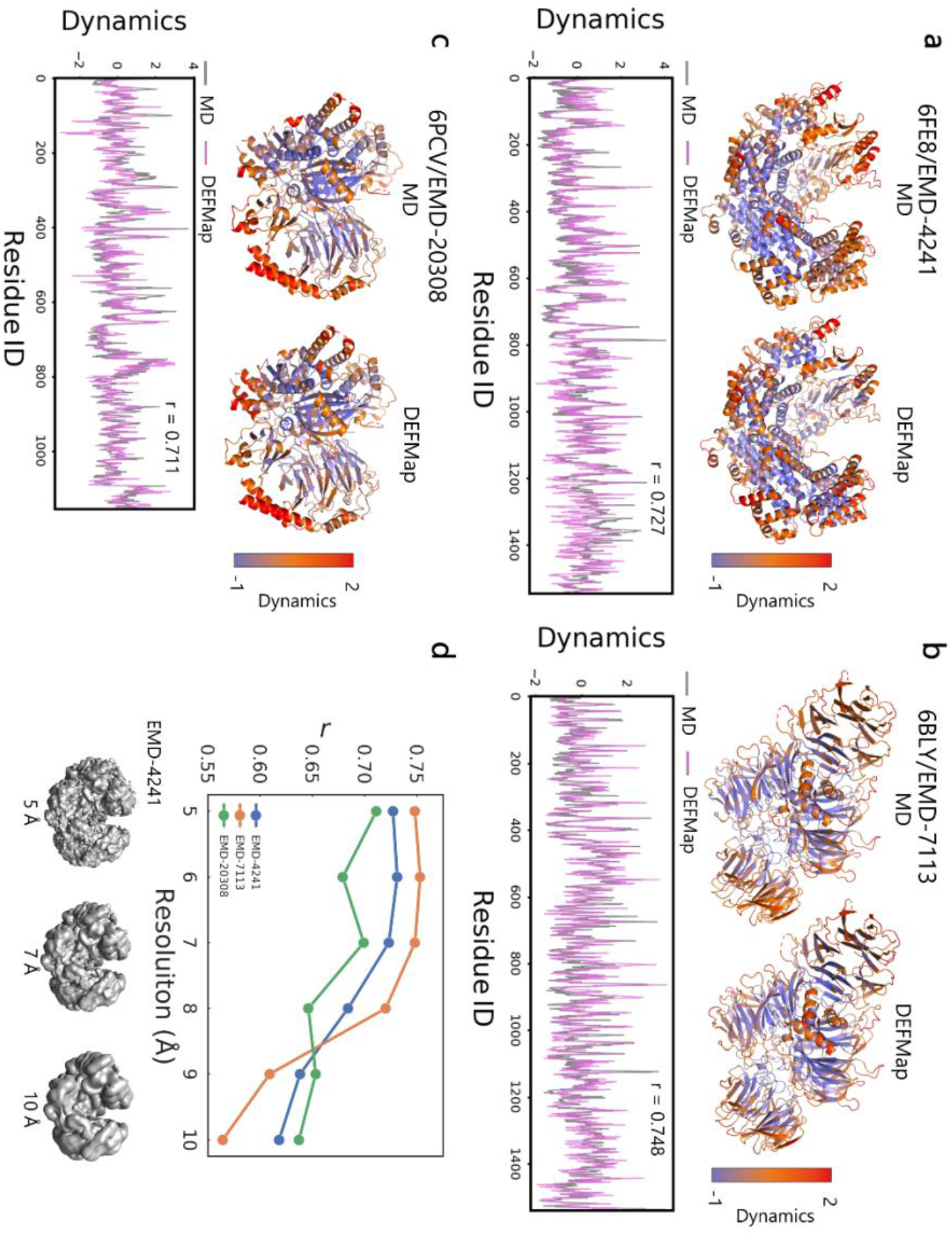
DEFMap performance for three proteins not included in the training dataset. **a-c,** Dynamicsmd and Dynamicsdefmap outputs for EMD-4241/6FE8 (**a**), EMD-7113/6BLY (**b**), and EMD-20308/6PCV (**c**). The cryo-EM maps were preprocessed by a 5 Å low-pass filter using the program EMAN2^39^. The proportions of secondary structure contents (α-helix/β-strand/others) are 0.56/0.06/0.38 (**a**), 0.05/0.43/0.52 (**b**), and 0.30/0.27/0.43 (**c**). The residue-specific values were calculated by averaging the normalized atomic values over each residue. The resulting values were mapped onto 3D atomic models (upper panels) using different colors as indicated in the color bars. The 3D Dynamicsmd and Dynamicsdefmap maps are shown on the left- and right-hand sides, respectively, of each panel. The bottom panels show the Dynamicsmd (gray) and Dynamicsdefmap (pink) profiles as a function of the residue IDs, numbered according to their order in the corresponding PDB file; *r* denotes the correlation coefficient between Dynamicsmd and Dynamicsdefmap. **d,** Dependence of DEFMap performance on map resolution. The plot shows the correlation coefficients between Dynamicsmd and Dynamicsdefmap obtained for different map resolutions. The maps of the training and other three datasets were preprocessed to identical resolutions with low-pass filters (e.g., the extraction of the Dynamicsdefmap data from the 10.0 Å low-pass filtered test map was carried out using the model trained by the 10.0 Å maps). The low-pass filtered cryo-EM maps of EMD-4241 at 5, 7, and 10 Å are shown at the bottom of the panel.

Confirming the consistency of the DEFMap predictions with experimentally derived dynamics properties is important for assessing the potential of the present method. Under the appropriate conditions, the dynamics of large proteins (such as those targeted by cryo-EM) can also be experimentally determined by HDX-MS^7^; using this approach, the dynamics information at the peptide fragment level is obtained by monitoring the effects of deuterium incorporation into protein amide groups (Supplementary Figure 4). We compared the Dynamicsdefmap of EMD-20308 with the publicly available HDX-MS data^33^. The average Dynamicsdefmap data for each fragment detected in the HDX-MS experiments were well-correlated with the corresponding HDX rates (Dynamicshdx-ms, *r* = 0.743) and Dynamicsmd data (*r* = 0.791), confirming that DEFMap accurately captured the local dynamics and can provide insight equivalent to that derived from experimental approaches (Fig. 3).

**Fig. 3:**
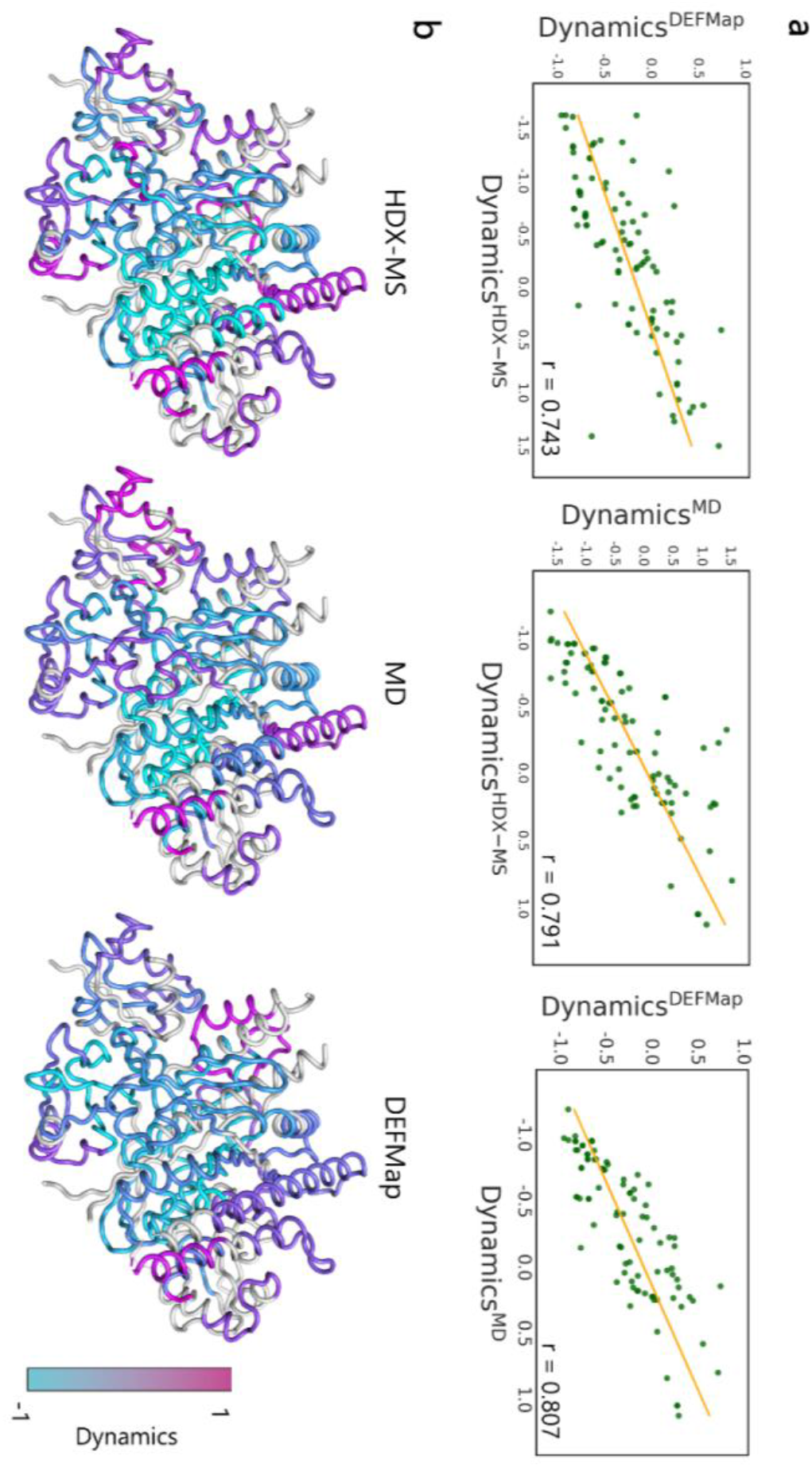
The relationship between DEFMap-determined and experimentally derived dynamics properties. **a,** A correlation plot of DEFMap-derived and HDX-MS-derived dynamics. The HDX exchange rates at 10^4^ s for Rac exchanger 1 complexed with G protein beta gamma subunits^33^ were normalized within the detected fragments and used as the experimental dynamics data. The residue-specific dynamics obtained from MD and DEFMap were converted to fragment-specific values by averaging them over each fragment in the atomic model. The HDX-MS *vs*. DEFMap (*r* = 0.743), HDX-MS *vs*. MD (*r* = 0.807), and MD *vs.* HDX-MS (*r* = 0.791) correlation plots are shown in the left, middle, and right panels, respectively, along with their corresponding regression lines (orange). **b,** The spatial distribution of the fragment-specific dynamics in the atomic models. The representative fragments-specific dynamics are mapped onto the atomic model using different colors as indicated in the color bar (HDX-MS, left; MD, middle; DEFMap, right). The fragments not detected in the HDX-MS experiments are colored gray.

Finally, we explored the potential of DEFMap for identifying molecular binding sites and investigating allosteric effects triggered by ligand binding. Ligand binding is a fundamental biological event and is often accompanied by the suppression of dynamics at the recognition interface. We therefore monitored dynamics changes associated with ligand-induced perturbations in the cryo-EM density maps. Particularly, we used DEFMap to detect ligand-induced dynamics changes for three pairs of apo/holo proteins not included in the training dataset (apo, holo: EMD-20080, EMD-20081^36^; EMD-9616, EMD-9622^37^; EMD-3957, EMD-3956^38^, Supplementary Table 4). We found good agreement between Dynamicsdefmap and Dynamicsmd profiles for the above pairs (Supplementary Figure 5). Moreover, the DEFMap-derived dynamics of the residues located near the binding partners were significantly suppressed by ligand binding (Fig. 4a, b and Supplementary Table 5), demonstrating that DEFMap could detect the conformational stabilization at the binding interface. Among the residues located at the interfaces (Supplementary Table 5), we observed significant dynamics suppression in the regions extensively discussed in the literature^36–38^. This finding suggests that DEFMap can identify the key interactions involved in complex formation using density data (Fig. 4b). We found additional dynamics suppression for regions distant from the ligand-binding site in the EMD-20080/EMD-20081 pair [Arabidopsis defective in meristem silencing 3 (DMS3)-RNA-directed DNA methylation 1 (RDM1) complex with defective RNA-directed DNA methylation 1 (DRD1) peptide]^36^ (Fig. 4c). In the apo state, DMS3 dimers establish contacts with a RDM1 dimer through their coiled-coil arms (middle panel in Fig. 4c). In the holo conformation, the binding partner, DRD1 peptide, is recognized by the RDM1 dimer, the coiled-coil arms of a DMS3 dimer (dimer 1 in Fig. 4c), and a hinge domain of an opposite DMS3 dimer (dimer 2 in Fig. 4c). DEFMap-based dynamics analysis showed that the binding of the DRD1 peptide induced suppression of dynamics at the RDM1-DMS3 interface and at the hinge domains in DMS3 dimer 1 (Fig. 4c), indicating that ligand binding allosterically stabilizes these regions. Interestingly, the static models constructed from the cryo-EM maps showed no substantial differences between the apo and holo proteins in these regions (Supplementary Figure 6). This finding further highlights the potential of DEFMap to provide dynamics information that cannot be determined from static tertiary structures.

**Fig. 4:**
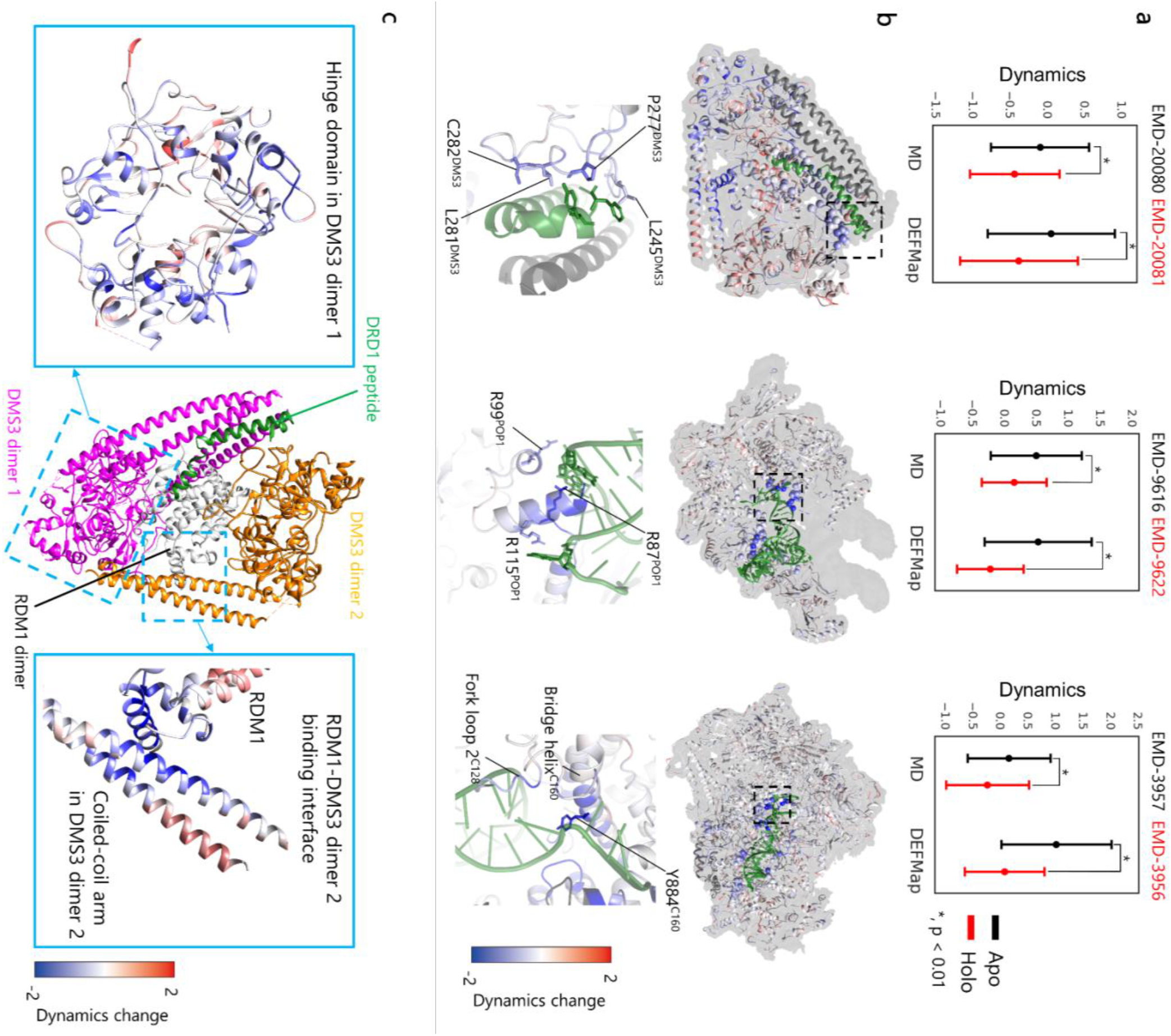
DEFMap-based detection of dynamics changes induced by ligand binding. **a,** Ligand-induced dynamics suppression at the binding interfaces. The dynamics at the binding interfaces in apo (black) and holo (red) proteins were calculated by averaging the Dynamicsmd and Dynamicsdefmap data for residues located within 5 Å of the binding partners, and the error bars indicate standard deviations (* symbols denote statistical significance at *p* < 0.01). **b,** The spatial distributions of ligand-induced dynamics changes derived from DEFMap. The dynamics changes were calculated by subtracting the Dynamicsdefmap of the apo state from those of the holo state and mapped onto the holo atomic models using different colors as indicated in the color bar (lower values denote regions with suppressed dynamics). The binding partners and disordered regions in the apo proteins are colored green and dark gray, respectively. In the overall view (upper panels), the Cα atoms of the residues located within 5 Å of the binding partners are represented as spheres, and the cryo-EM maps are colored gray. The lower panels show enlarged views of the regions indicated by dashed rectangles in the upper panels. The relevant residues extensively discussed in the literature are represented as sticks, with the subunit names indicated as superscripts. **c,** DEFMap detection of ligand-induced allosteric change. Enlarged views of the regions discussed in the main text are shown in the left- and right-hand side panels, respectively. The color bar represents the ligand-induced dynamics changes.

Cryo-EM SPA opened a new era in molecular biology by enabling researchers to solve 3D structures of macromolecules and supramolecules at high resolution. In this study, we showed that a deep neural network-based approach, DEFMap, has successfully extracted “hidden” dynamics information only from static 3D cryo-EM density maps. The DEFMap approach is not restricted by the molecular sizes and the complexity of the systems because DEFMap infers dynamics from local density data, as shown in the above results. We expect that complementary use of DEFMap with conventional cryo-EM SPA will accelerate the elucidation of the molecular mechanisms underlying biological events. Furthermore, DEFMap will help researchers readily access the dynamics properties of biological molecules, as this open-source tool does not require additional expensive and/or time-consuming experiments or in-depth expertise. We believe that in the future, DEFMap will accelerate data-driven structural investigations aiming to understand protein functions and to develop strategies for molecular targeted therapy against various diseases. This study bridges experimental data, deep learning approaches, and MD simulations, allowing for accurate extraction of hidden information, which humans could not utilize, from the data. It is expected that this advanced strategy would open a new multidisciplinary science area which integrate experimental science, simulation science, and data science.

## Methods

### Molecular dynamics simulations

All-atom MD simulations for macromolecules in the present datasets (34 molecules) are carried out to obtain RMSF values that used for training of a deep learning model in DEFMap or evaluation of the performance. The selected macromolecules included proteins or the complex with DNA/RNA which are conveniently handled in the simulations. The initial coordinates used in the simulations were obtained from the PDB (https://www.rcsb.org) and processed using the Structure Preparation module of the Molecular Operating Environment (MOE) software (Chemical Computing Group, Montreal, Canada). Briefly, loops were modeled for disordered regions containing less than seven residues, and other non-natural N-termini and C-termini were capped with acetyl and formyl groups, respectively. The addition of hydrogen atoms and the generation of topology files were carried out using the pdb2gmx module in the GROMACS package^40^. All MD simulations were carried out with periodic boundary conditions (PBC), using GROMACS^40^ on an NVIDIA GeForce GTX 1080 GPU. The shape of the periodic cell was octahedron. The Amber ff99SB-ILDN force field was used for proteins, nucleotides, and ions^41^, while the TIP3P potential was used for water^42^. Water molecules were placed around the substrate model with an encompassing distance of 10 Å, and counterions (NaCl) were added to neutralize the system. Electrostatic interactions were calculated using the particle mesh Ewald (PME) method^43^ with a cutoff radius of 10 Å, and a nonbonded cutoff of 10 Å was applied for van der Waals interactions. The P-LINCS algorithm was used to constrain all bond lengths to their equilibrium values^44^. After energy minimization of the fully solvated models, the resulting systems were equilibrated for 100 ps under constant number of molecules, volume, and temperature (NVT) conditions, followed by a 100 ps run under constant number of molecules, pressure, and temperature (NPT) conditions, with the heavy atoms of the macromolecules held in fixed positions. The temperature was maintained at 298 K by velocity rescaling with a stochastic term^45^, while Parrinello-Rahman pressure coupling^46^ was used to keep the pressure at 1 bar, with the temperature and pressure time constants set to 0.1 and 2 ps, respectively. Subsequently, production runs of 20 ns were carried out under NPT conditions without positional restraints. After PBC corrections, the generated trajectories were aligned using overall Cα atoms, followed by the calculation of the RMSF values (Å) for heavy atoms using the rmsf module of GROMACS. The logarithmic RMSFs were then used to represent dynamics properties.

### Data preparation for DEFMap

Twenty-five cryo-EM density maps were selected and downloaded from the EMDB/PDB (Supplementary Table 1, average overall resolution = 3.62 ± 0.46) to train the 3D-CNN model. Their resolutions are relatively high compared to the average overall resolution of cryo-EM maps deposited in 2019 (5.6 Å). To evaluate the DEFMap potential for dynamics analysis, the additional nine cryo-EM density maps were selected and downloaded from the EMDB/PDB (Supplementary Tables 3 and 4). The maps were rescaled to 1.5 Å/pixel and low-pass filtered using EMAN2.3^39^. Subsequently, the intensities were normalized within each map, and any negative values were discarded. Each grid point in the maps was associated with the MD-derived logarithmic RMSF of the nearest atom in the voxelized coordinates. The resulting maps were sub-voxelized to generate the input density data, with 10^3^ voxels distributed over 15^3^ Å^3^. The training data were augmented by x-, y-, and z-axis rotations of 90°, 180°, and 270°. The preprocessed maps with an overall resolution of 5 Å were mainly used in this study unless otherwise stated. Voxelization of the atomic models was carried out by high-throughput molecular dynamics (HTMD)^47^. All preprocessing procedures were performed using Python.

### Construction of deep neural networks

The architecture of the neural network used in DEFMap included 3D convolutional blocks and dense blocks. The 3D convolutional blocks consisted of three, 3D convolutional layers with leaky ReLU activation, max pooling, and dropout. The kernel size for convolution, the maximum pooling size, and the dropout ratio were set to 4 × 4 × 4, 2 × 2, and 0.2, respectively. Different filter sizes (64, 128, and 256) were applied to the three 3D convolutional layers. The dense blocks consisted of two dense layers of 1,024 units with leaky ReLU activation and a dense layer of a unit with identity activation. The epochs, learning rate, and batch size hyperparameters were set to 50, 0.00005, and 128, respectively, and early stopping with a patience interval of three epochs was utilized to avoid overfitting. All models were trained using an NVIDIA Tesla V100 GPU with 16 GB of memory. The Keras^48^ library with Tensorflow backend was employed in the calculations.

### Postprocessing and visualization of output from the neural networks

Postprocessing of the atomic dynamics values calculated by DEFMap for the subsequent validation and analyses was carried out as follows: the output values were normalized and then averaged over each residue; the residue-specific values (Dynamicsdefmap) were then assigned to the atomic models as temperature factors with HTMD^47^ and visualized using the PyMOL^49^ and UCSF Chimera^50^ programs. All postprocessing procedures were performed using Python.

## Code availability

The Python code used to implement DEFMap is available on GitHub at https://github.com/clinfo/DEFMap. DEFMap is licensed under MIT licence.

## Data availability

All datasets used in this paper are publicly available via the PDB (https://www.rcsb.org/). Detailed descriptions of the datasets are provided in Supplementary Tables 1, 2, and 3.

## Acknowledgements

This work was supported by MEXT as “Priority Issue on Post K computer (Building Innovative Drug Discovery Infrastructure Through Functional Control of Biomolecular Systems)” and JSPS KAKENHI Grant Number JP17K15106 to SM.

## Author contributions

S.M., S.I., K.T., and Y.O. designed the study. S.M., and S.I. performed the pre-processing of cryo-EM maps, design the neural networks, and carried out model training and DEFMap analyses. S.M. and M.A. performed the MD calculations. S.M. and T.K. prepared the original cryo-EM map data. K.T. and Y.O. conceived the project. S.M. and S.I. wrote the manuscript. All authors discussed the research, edited the manuscript, and approved its final version.

