## Supplementary Information for "Extraction of Protein Dynamics Information Hidden in Cryogenic Electron Microscopy Maps Using Deep Learning"

[1] Medical Sciences Innovation Hub Program, RIKEN Cluster for Science, Technology and Innovation Hub, 1-7-22 Suehiro-cho,Tsurumi-ku, 230-0045 Kanagawa, Japan

[2] Graduate School of Pharmaceutical Sciences, Kyoto University, 46-29 Yoshidashimo-Adachicho, Sakyo-ku, 606-8501 Kyoto, Japan.

[3] Graduate School of Medicine, Kyoto University, 53 Shogoin-Kawaharacho, Sakyo-ku, 606-8507 Kyoto, Japan.

[4] Institute for Protein Research, Osaka University, 3-2 Yamadaoka, Suita, 565-0871 Osaka, Japan.

[5] RIKEN Center for Advanced Intelligence Project, 1-4-1 Nihonbashi, Tokyo 103-0027, Japan.

### Author information
[†]These authors contributed equally to this study.

### Corresponding authors

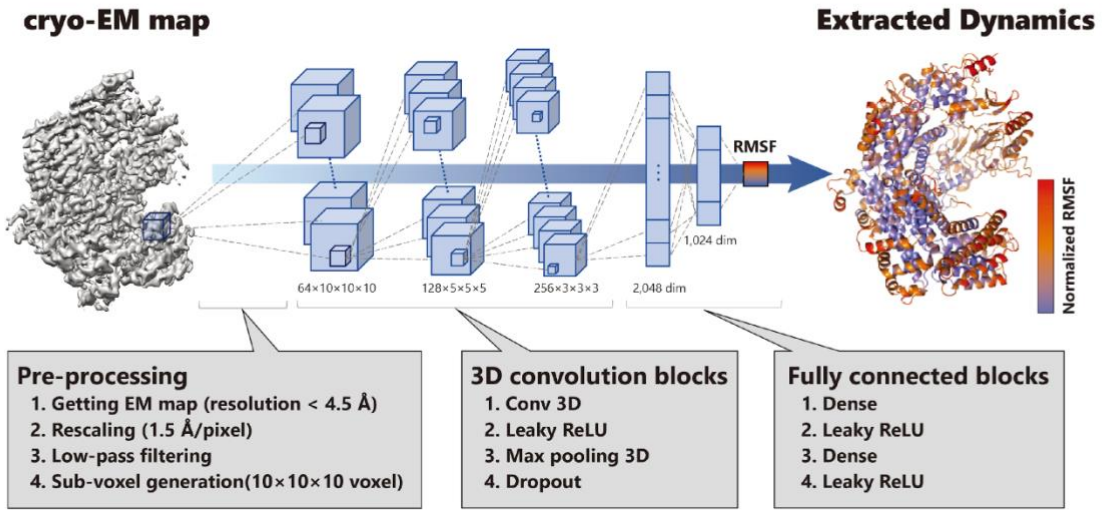


**Supplementary Figure 1: Details of pre-processing and 3D-CNN model of DEFMap.** The DEFMap system is composed of three steps: preprocessing, neural network processing, and visualization of dynamic properties after postprocessing (see Methods section). In the preprocessing step, 10×10×10 sub-voxels ($15^3$ Å$^3$) are generated from the original cryo-EM map through low-pass filtering and rescaling of the grid length to 1.5 Å/pixel. In the neural network processing step, each generated voxel is converted to the corresponding logarithmic RMSF value, using 3D-CNN and fully connected neural networks. In the visualization step, the calculated RMSF values are averaged over each residue after normalization and then mapped onto the corresponding atomic models. The different colors represent the normalized logarithmic RMSFs.

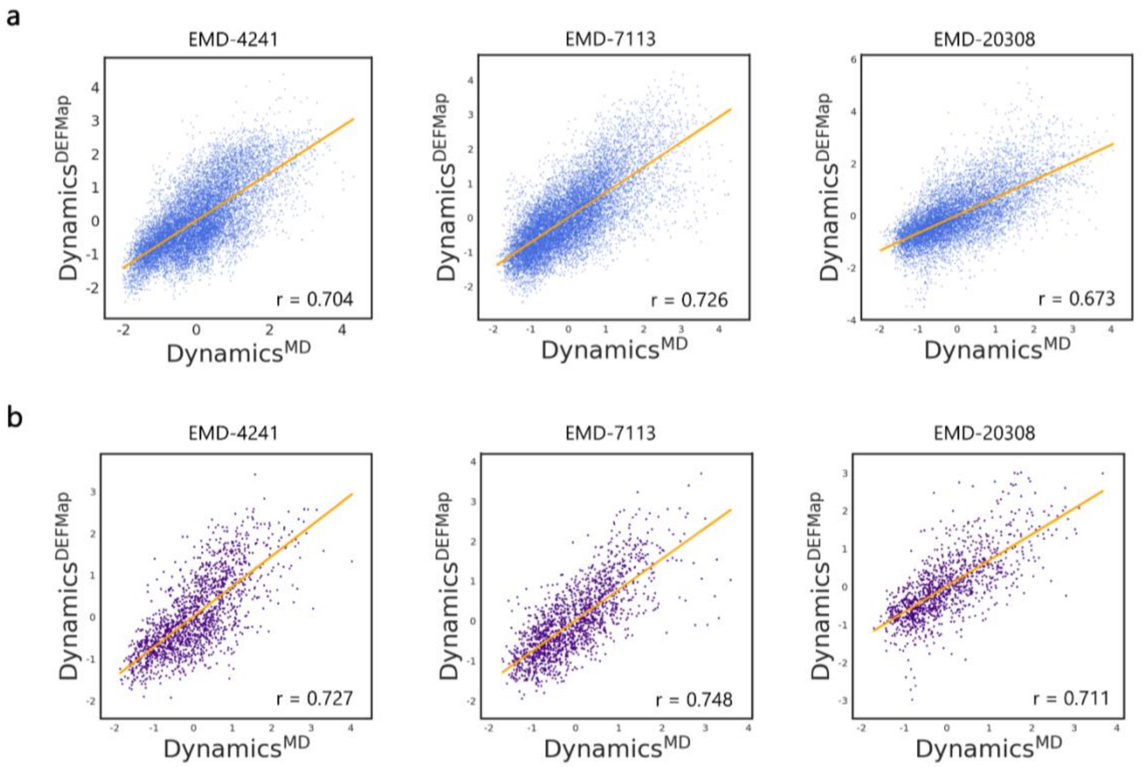


**Supplementary Figure 2. Correlation plots for MD-derived and DEFMap-determined dynamics for proteins not included in the training dataset. a, b,** Normalized atomic (**a**, blue) and residue-averaged (**b**, purple) values are shown in the correlation plots; the regression lines are colored orange; *r* denotes the correlation coefficient.

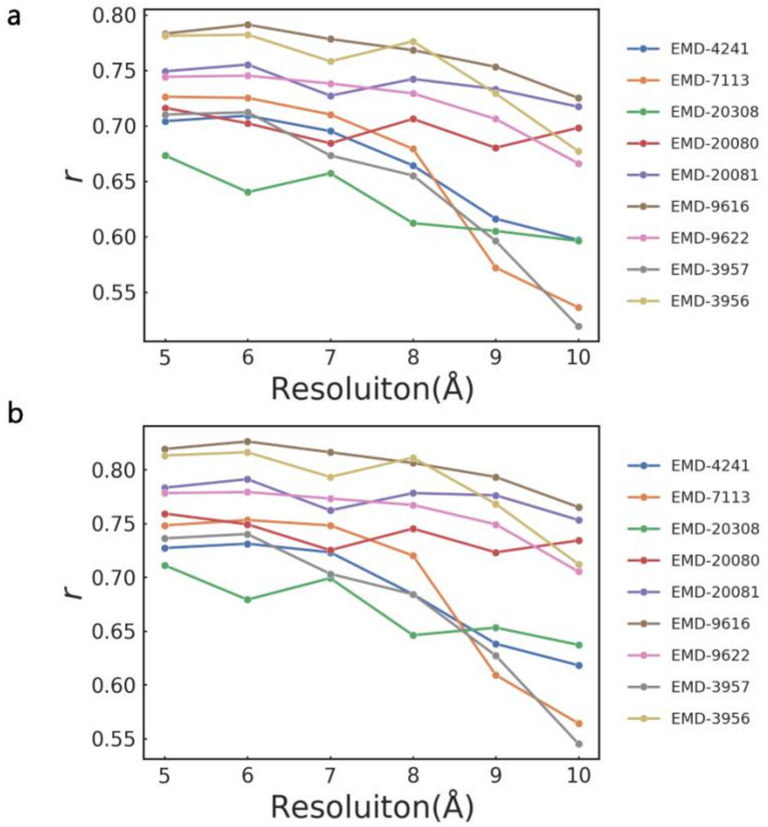


**Supplementary Figure 3. The map resolution dependence of the dynamics extraction accuracy for test macromolecules (Supplementary Tables 3 and 4). a, b,** Correlation coefficient values between the MD- and DEFMap-derived atomic-specific (**a**) and residue-specific (**b**) dynamics at different resolutions. The cryo-EM maps of the training and the evaluation datasets were preprocessed to identical resolutions with low-pass filters.

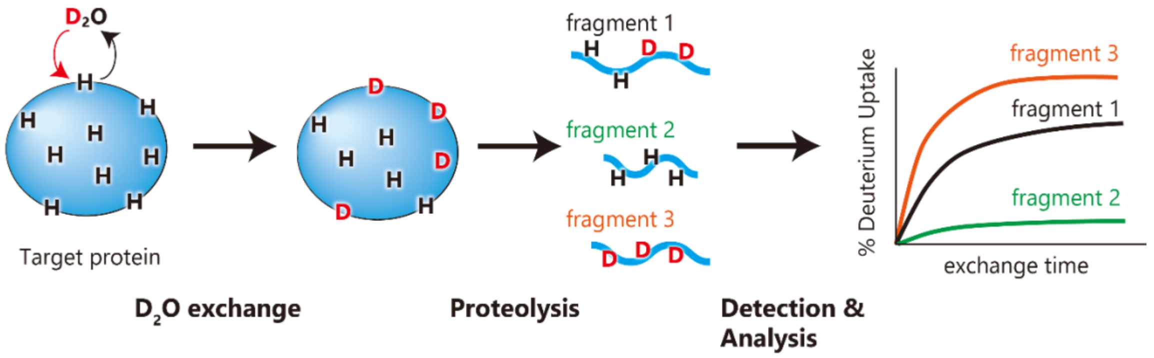


**Supplementary Figure 4. Overview of HDX-MS experiments.** The HDX-MS method determines local dynamics at the peptide fragment level by monitoring deuterium incorporation from the $D_2O$ solvent into protein amide groups. The extent of the incorporation depends on the solvent accessibility and the local chemical environment of the exchangeable protons. After exchange reactions during a certain incubation time and quenching, the labeled protein is proteolyzed to generate a series of peptide fragments using proteases. The deuterium incorporation ratio for each peptide fragment are determined by measuring the mass shifts by mass spectrometry.

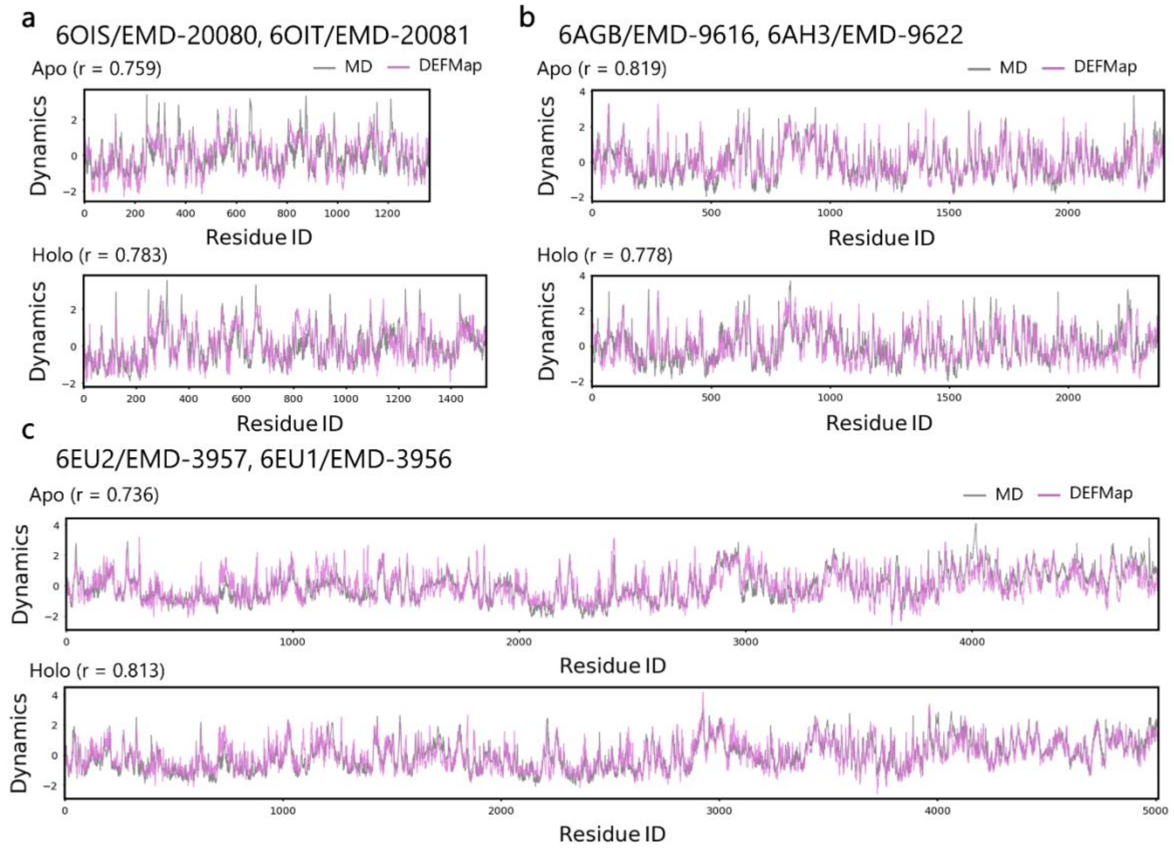


**Supplementary Figure 5. Residue-specific dynamics profiles for apo/holo proteins. a, b,** $Dynamics_{MD}$ (grey) and $Dynamics_{DEFMap}$ (pink) datasets for 6OIS/EMD-20080 and 6OIT/EMD-20081 (**a**), 6AGB/EMD-9616 and 6AH3/EMD-9622 (**b**), and 6EU2/EMD-3957 and 6EU1/EMD-3956 (**c**), respectively, plotted against the residue IDs, numbered according to their order in the coordinate files. *r* denotes the correlation coefficient between $Dynamics_{MD}$ and $Dynamics_{DEFMap}$.

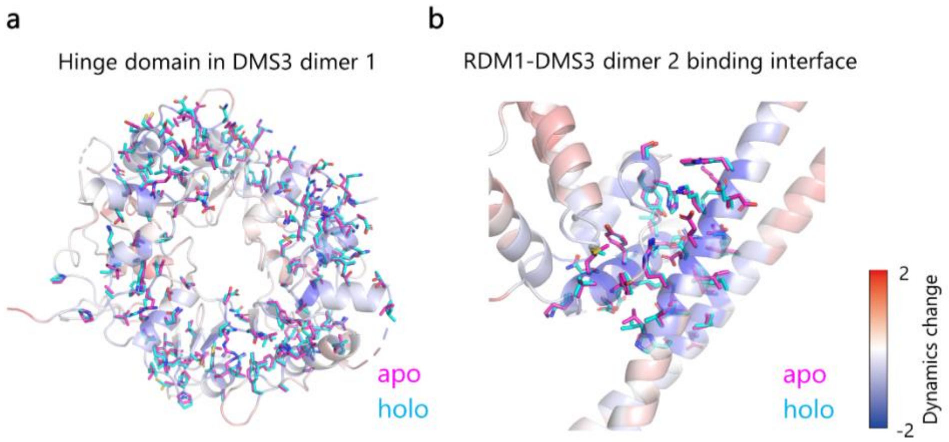


**Supplementary Figure 6. Atomic models of residues showing allosteric dynamics changes induced by ligand binding in the DMS3-RDM1 complex with DRD1 peptide.** **a, b,** The main chains of the hinge domain in DMS3 dimer 1 (**a**) and in the RDM1-DMS3 dimer 2 binding interface (**b**) are colored according to the extent of ligand-induced changes in dynamics, following the same notation used in Fig. 4b. The side chains of residues showing dynamics values of less than -0.5 are represented as sticks.

**Supplementary Table 1. List of EMDB/PDB IDs of molecules used in model training.**

| EMDB ID | PDB ID | Resolution (Å) |
|---|---|---|
| EMD-9590 | 6ACF | 3.00 |
| EMD-0010 | 6GJ3 | 4.30 |
| EMD-9623 | 6AHC | 3.45 |
| EMD-7303 | 6BX3 | 4.30 |
| EMD-9588 | 6ACC | 3.60 |
| EMD-3824 | 5OJS | 3.70 |
| EMD-20523 | 6PXW | 3.10 |
| EMD-3984 | 6EZ8 | 4.00 |
| EMD-7436 | 6C9I | 3.09 |
| EMD-7465 | 6CET | 4.40 |
| EMD-7062 | 6B70 | 3.70 |
| EMD-8653 | 5VAI | 4.10 |
| EMD-3460 | 5MBV | 3.80 |
| EMD-3583 | 5MZ6 | 3.80 |
| EMD-6630 | 3JCZ | 3.26 |
| EMD-8191 | 5K0Z | 2.80 |
| EMD-6324 | 3JA7 | 3.60 |
| EMD-6940 | 5ZQZ | 4.20 |
| EMD-9657 | 6IFU | 3.05 |
| EMD-3601 | 5N8O | 3.90 |
| EMD-9887 | 6KUJ | 3.40 |
| EMD-20457 | 6PR5 | 3.30 |
| EMD-0065 | 6GTG | 3.27 |
| EMD-9627 | 6AHU | 3.66 |
| EMD-9900 | 6K0B | 4.30 |

**Supplementary Table 2. Correlation coefficients of MD-derived dynamics with local cryo-EM map intensities and DEFMap-derived values.**

| Fold # | Cryo-EM Map | DEFMap | Test Protein (EMDB ID) | Number of Voxels |
|---|---|---|---|---|
| 1 | 0.382 | 0.742 | EMD-3984 | 18767 |
| 2 | 0.572 | 0.821 | EMD-6324 | 37752 |
| 3 | 0.331 | 0.529 | EMD-9623 | 41995 |
| 4 | 0.067 | 0.300 | EMD-7303 | 9543 |
| 5 | 0.526 | 0.523 | EMD-0010 | 15909 |
| 6 | 0.546 | 0.750 | EMD-9657 | 15916 |
| 7 | 0.474 | 0.707 | EMD-3824 | 25927 |
| 8 | 0.563 | 0.733 | EMD-9588 | 22741 |
| 9 | 0.696 | 0.898 | EMD-9590 | 20950 |
| 10 | 0.670 | 0.769 | EMD-7436 | 21581 |
| 11 | 0.335 | 0.747 | EMD-3583 | 7704 |
| 12 | 0.637 | 0.806 | EMD-0065 | 9712 |
| 13 | 0.430 | 0.606 | EMD-7062 | 20200 |
| 14 | 0.461 | 0.657 | EMD-9627 | 18916 |
| 15 | 0.626 | 0.744 | EMD-9887 | 10705 |
| 16 | 0.747 | 0.740 | EMD-6630 | 21321 |
| 17 | 0.147 | 0.510 | EMD-8653 | 8266 |
| 18 | 0.209 | 0.547 | EMD-7465 | 12808 |
| 19 | 0.349 | 0.709 | EMD-6940 | 15703 |
| 20 | 0.455 | 0.519 | EMD-9900 | 10369 |
| 21 | 0.609 | 0.641 | EMD-3460 | 22068 |
| 22 | 0.465 | 0.685 | EMD-20457 | 6992 |
| 23 | 0.449 | 0.655 | EMD-20523 | 10184 |
| 24 | 0.479 | 0.634 | EMD-3601 | 10024 |
| 25 | 0.374 | 0.651 | EMD-8191 | 8877 |
| **Average ± SD** | **0.464 ± 0.164** | **0.665 ± 0.124** | | |

**Supplementary Table 3. List of EMDB/PDB IDs of molecules to evaluate the DEFMap potential toward dynamics analysis against the proteins not included in the training dataset.**

| EMDB ID | PDB ID | Resolution (Å) |
|---|---|---|
| EMD-4241 | 6FE8 | 4.10 |
| EMD-7113 | 6BLY | 3.36 |
| EMD-20308 | 6PCV | 3.20 |

**Supplementary Table 4. List of EMDB/PDB IDs for apo/holo pair molecules to evaluate the potential of DEFMap in structural biology studies.**

| EMDB ID | PDB ID | State | Resolution (Å) |
|---|---|---|---|
| EMD-20080 | 6OIS | apo | 3.60 |
| EMD-20081 | 6OIT | holo | 3.50 |
| EMD-9616 | 6AGB | apo | 3.48 |
| EMD-9622 | 6AH3 | holo | 3.48 |
| EMD-3957 | 6EU2 | apo | 3.40 |
| EMD-3956 | 6EU1 | holo | 3.40 |

**Supplementary Table 5. List of residues located within 5 Å of the binding partners in apo/holo pairs.**

| EMDB/PDB ID | Residues |
|---|---|
| EMD-20080/6OIS<br>EMD-20081/6OIT | (A)[a] R64, H118, T120, K121, A124, L128 (B) T62, N63, R64, T120, K121, A124, L128 (D) Q239, D244, L245, Q246, R247, D250, D276, P277, A278, L280, L281, C282, S285, Y286, G287, Y288 (E) R55 |
| EMD-9616/6AGB<br>EMD-9622/6AH3 | (B) A83, S84, S85, T86, R87, I88, Q90, R99, S102, H103, R115, R118, E119, K122, S123, D124 (D) Q205, Y255, R256, D259, R263, K264, K266, S267 (E) V2, R3, L4, S70, L71, Q73, R87 (J) H49, K52, N55, P91, S92, K93, G94, Q95, S96, L97, S98, K99, K115, T118, L119, N123, L124, D125, T139, F140, L141, K142, K144, S145, R152, R182, R239, K244 (K) K75, K76, K78 |
| EMD-3957/6EU2<br>EMD-3956/6EU1 | (A) K150, R152, V164, K165, K166, W185, V186, K188, K189, R372, R378, Q477, P478, T875, A876, T879, A880, G883, Y884, R887, S1133, K1134, V1135, K1216, R1373, F1374, E1390, K1391, T1392, D1394 (B) R227, F477, E478, K479, R481, V483, K1034, G1053, R1054, R1056, L1060, R1061, G1063, E1064, M1065 (E) G89, V90, T117, P118, S119 (O) S233, D234, L235 |

[a] Uppercase letters in parentheses denote chain names in the coordinate files.
